# Parasite transmission in size-structured populations

**DOI:** 10.1101/2022.12.07.519457

**Authors:** Kelsey E Shaw, Rebecca E Cloud, Raeyan Syed, David J Civitello

## Abstract

Host heterogeneity can impact parasite transmission, but determining underlying traits and incorporating them into transmission models remains challenging. Body size is easily measured and influences numerous ecological interactions, including transmission. In the snail-schistosome system, larger snails have a higher exposure to parasites but lower susceptibility to infection per parasite. We quantified the impact of size-based heterogeneity on population-level transmission by conducting transmission trials in differently size-structured snail populations and competing size-dependent transmission models. Populations with greater proportions of large snails had lower prevalence, and small snails were shielded from infection by co-occurring large conspecifics. Using the winning size-explicit model, we then estimated that schistosome transmission varies dramatically across time due to seasonal changes in snail population size structure. Thus, incorporating traits such as body size, which are impacted by and directly affect host ecology, into transmission models could yield insights for natural dynamics and disease mitigation in many systems.

**Open Research Statement:** All experimental data and novel code used for data analysis are currently available on Figshare via this private link: https://figshare.com/s/7d70b88220b912e7eec5. Data will be made public on Figshare upon acceptance. Data set utilized for Figure 4 are from the following source: Rumi, A., D. Gutiérrez Gregoric, and A. Roche. 2009. Tendencias Del Crecimiento Individual en Poblaciones Naturales de Biomphalaria spp. (Gastropoda, Planorbidae) en la Cuenca del Plata, Argentina. *Comunicaciones de la Sociedad Malacológica del Uruguay*. URL: https://www.redalyc.org/articulo.oa?id=52414008002

## I. Introduction

Parasite transmission drives the ecological and evolutionary dynamics of host-parasite interactions. Therefore, demystifying the host and parasite factors that influence transmission is critical for characterizing disease dynamics and successfully managing any host-parasite system. In particular, transmission theory that excludes host variation and instead treats hosts as uniform individuals with identical infection risk matches data poorly and cannot address critical themes in disease ecology, such as superspreading, parasite aggregation, and host contact networks (Lloyd-Smith et al. 2005, Van der Waal and Ezenwa 2016). For example, early in the Sars-CoV-2 pandemic, models that did not include variability in traits like infectiousness or contacts often overestimated values for R_0_, with implications for public health recommendations (Gomes et al. 2022). In order to more accurately predict disease dynamics in natural populations, it is important to consider the sources of host variation and their consequences for transmission because hosts can differ in many ways (Dwyer et al. 1997, Hall et al. 2007, Strauss et al. 2019).

Body size is a one host trait that can be highly heterogenous and may impact parasite transmission in many systems. For example, larger individuals may provide a greater target area for vectors (Poulin 2004, Yan et al. 2021), may provide more resources and thus tolerate a larger burden of macroparasites (Hechinger et al. 2019), may be more apparent to parasites that actively seek out their host using chemical cues (Théron et al. 1998, Mayer et al. 2021), or may encounter a greater abundance and diversity of parasites if they are exploring larger territories(Han et al. 2015). Furthermore, body size has the potential to vary drastically among individuals in a population due to age, genetics, feeding history, and/or environmental stressors (Kooijman, S.A.L.M. 2010). The snail-schistosome system is an excellent study system for investigating the impact of host variation in body size on parasite transmission.

Schistosomes are the causative agent of schistosomiasis, a Neglected Tropical Disease that is second only to malaria in its worldwide impact on human health and disproportionately affects individuals in poor, rural environments (Gryseels et al. 2006). Over 200 million people worldwide are currently infected, and schistosomiasis accounts for approximately 200,000 deaths per year (World Health Organization 2022). Schistosomes have a complex life cycle and obligately cycle between mammalian and snail hosts. One source of host heterogeneity that impacts schistosome transmission is snail body size: larger snails experience higher exposure rates but are less likely to become successfully infected on a per parasite basis (Niemann and Lewis 1990, Théron et al. 1998). Snail body size can be influenced by many factors such as age, genetics, and resource availability, but this relationship between body size, exposure, and susceptibility has been demonstrated to occur regardless of the underlying size-determining mechanism (Richards et al. 1992, Larson et al. 2014). The importance of body size in transmission to the snail host becomes even more apparent when considering the vast range in host snail body size. For example, a neonate *Biomphalaria glabrata*, an intermediate host species for *Schistosoma mansoni*, is born at ∼0.75 mm and can grow up to 40 mm under natural conditions over its lifetime (Jarne et al. 2011). This growth trajectory corresponds to a ∼150,000-fold increase in biomass, akin to the difference between a mouse and an elephant (White 2010).

Despite the recognition that body size influences these mechanistic components of schistosome transmission decades ago in isolated, individual snails, the consequences of this relationship when scaled to size-structured populations have remained uninvestigated. Schistosome exposure in a host is irreversible, meaning a schistosome cannot leave a snail once it has entered, even if it fails to successfully infect the host. Therefore, snails that are invaded by parasites indirectly reduce the infection risk of others in the population (King et al. 2011). If size-dependent transmission effects scale up to the population level, then we predict that the presence of larger hosts will contribute to a within-species dilution effect by shielding their smaller conspecifics from parasite exposure while also being less likely to become infected themselves.

We investigated this hypothesis by conducting laboratory transmission trials of *Schistosoma mansoni* in *Biomphalaria glabrata* populations of varying size structures. We then used these data to compete candidate models of size-dependent transmission, and additionally used existing field data of *B. glabrata* population size structures to estimate how parasite success may change under natural conditions over time solely due to the dynamics of population size structure. In many host-parasite systems researchers are still in the nascent phase of breaking down the complex interaction of host traits that result in heterogeneous parasite transmission (Sweeny and Albery 2022). In the snail-schistosome system, body size is a measurable trait with the potential to influence parasite transmission via multiple mechanisms, but it remains relatively unexplored how this individual-level trait scales up to influence transmission within and between populations. Incorporating this fundamental host trait into our understanding of schistosome transmission could yield important insights into transmission risk and disease control of this widespread and debilitating parasite, and this approach can be applied broadly to other disease systems and critical traits.

## II. Methods

### Snail Maintenance

*B. glabrata* snails of the NMRI strain were maintained under favorable conditions. Snails were kept in HHCOMBO artificial lake water (Baer and Goulden 1998) at 26°C with a 12:12 light:dark cycle. Snails were fed a diet of fish flakes (Omega One) and chicken feed (Nutrena Meatbird Crumbles) suspended in 1% agar *ad libitum*.

### Experimental Design

We conducted the experiment using a fully factorial design: five population size-structures, each with a total of 18 snails, were exposed to three densities of parasites in 24-hour transmission trials in 15-liter tanks, creating 15 total treatment combinations. *B. glabrata* were first assigned into three size classes by shell diameter: “small” (2-3 mm), “medium” (6-8 mm), and “large” (12-15 mm). Different size-structures were determined by the ratio of snails from each of the three size classes. The uniform size structure consisted of 18 snails from the same size class: “uniform small” (1:0:0), “uniform medium” (0:1:0), and “uniform large” (0:0:1). “Equal” size-structured mesocosms had 6 snails from each size class (1:1:1). “Small skewed” size structures contained 12 snails from the smallest size class and 3 snails from both the medium and large size classes (4:1:1). Snail size classes and population size structures were chosen to encompass the range of size variability observed in nature (see Fig 4) and also to span a sufficient gradient to fit and evaluate size-dependent transmission models. Parasite densities also varied between the tanks: 36 (2 miracidia/snail), 144 (8 miracidia/snail), and 256 parasites per tank (14 miracidia/snail). We obtained *S. mansoni* eggs from experimentally infected mice livers and purified and hatched eggs to obtain miracidia (Dinguirard et al. 2018). We conducted four randomized blocks of these trials in which each block contained one replicate of each treatment. To control for potential batch effects on parasite infectivity across the temporal blocks, a control group consisting of all medium snails individually placed in 24 well plates was simultaneously exposed to each of the parasite densities.

To quantify size-based susceptibility we also conducted transmission experiments in which 120 snails ranging from 0.5mm to 15mm were individually placed in well plates with 5 mL COMBO. These snails were exposed to either 2 or 6 miracidia over 24 hours. Due to the small volume of water used in the individual well plate exposures, we assume that snails were exposed to all parasites present in these trials, an assumption justified by prior experiments (Niemann and Lewis 1990).

### Infection Diagnosis

24 hours post-exposure, all snails were collected, sorted by size and treatment group, and maintained for 5 weeks (longer than the minimal prepatent period for *S. mansoni* in favorable conditions). Snails were diagnosed visually by parasite shedding in individual well plates at 4- and 5-weeks post-exposure to obtain prevalence data (Asch 1972). Snails were diagnosed as “infected” if we observed *S. mansoni* cercariae in their well plate, or “uninfected” if no cercariae were detected. Snails infected at week 4 were sacrificed, while uninfected snails were returned to their tanks to repeat this process at week 5. Snails that died while prepatent or between weeks 4 and 5 of the shedding process were not included in our results (6.25% of all snails).

To compare prevalence data amongst the various size structures and size classes, we used the glmmTMB package in R (Brooks et al. 2017) to create a generalized linear mixed model (GLMM) accounting for the random effects between individual tanks and exposure dates (bionomial probability distribution with a logit-link function). We then used the emmeans package to conduct an estimated marginal means (least-squares means) *post hoc* test (Lenth 2022).

### Model Creation and Maximum Likelihood Estimation of Parameters

We then built deterministic size-dependent transmission models for parameterization and competition using the experimental data (Box 1). The models used coupled ordinary differential equations to track the densities of susceptible snails (S_i_) and infected snails (I_i_) in each size class (i) as well as free living parasites in the water (P). In our experiments, snails and parasites were all added at t_0_ and hosts were removed 24 hours later at t_1_, therefore we only focus on a single timestep and do not include processes on a longer time scale such as host birth and death. Each model separates the transmission process into two underlying processes: exposure to parasites and susceptibility to infection given exposure(Civitello and Rohr 2014). The models varied based on whether these processes were functions of snail body size (represented as the longest shell diameter, hereafter “length”). In the null model, all functions of length are constants, therefore no variation in prevalence by size-structure can be predicted. We also considered two models in which one parameter varied with body size: (1) the size-dependent exposure-only, in which only exposure is a function of host body size, and (2) the size-dependent susceptibility only, in which only susceptibility is a function of body size. Finally, we considered a fully size-dependent model, which varies both exposure (ε) and susceptibility (σ) with host body size. All models yield an analytical solution for snail infection prevalence at time t_1_ (Box 1). The three models that contain some element of size-dependence all predict that different size classes within a single size-structured population can experience different levels of infection prevalence and that populations of differing size-structures can vary in their overall population infection prevalence. For all models with a size-dependent component, we investigated two alternative parsimonious representations: linear and exponential functions of host length (Appendix S1: Figure S1).

For each model, we solved for the expected prevalence in each size class present analytically, given the size structure and parasite density of that treatment (Box 1). We then used the bbmle package in R to conduct maximum likelihood estimation to estimate parameters for ε and σ for each model using the binomial error distribution with the probability of “success”, i.e., infection, equal to the expected prevalence for that size class given the experimental conditions and parameter values (Allen, Linda 2010, Strauss et al. 2019, Bolker and R Development Core Team 2021). This probability distribution provides the likelihood for our observations of infection status for snail hosts in each size class, which we combined to generate an overall likelihood for all observations. Lastly, to control for potential differences in infectivity among parasites across the different temporal blocks, we incorporated a block effect for the baseline susceptibility parameter, σ_0_. Using corrected Akaike’s information criteria (AICc) in R (R Core Team 2021), we competed the fully size-dependent model against the size-dependent exposure only, the size-dependent susceptibility only, and the size-independent null model.

### Application to field data

We then estimated the influence of population size structure on schistosome transmission for a snail population in an endemic setting experiencing natural variation in size structure throughout the year. We used WebPlotDigitizer (Rohatgi 2020) to gather snail size data over time from a population of *Biomphalaria tenagophilia* in del Plata Basin, Argentina from 2000-2001 (Rumi et al. 2009). WebPlotDigitizer is a free, open source, web-based platform that allows for measurement and quantification of images. We took the histogram plots of snail sizes across time in Figure 1 of Rumi et al. 2009 and imported them into WebPlotDigitizer. Then, we labeled the corresponding value of the X axis and the Y axis in the original figure and marked the data points. After that, WebPlotDigitizer can automatically extract the specific values of the marked data points. The code of WebPlotDigitizer is available at https://github.com/ankitrohatgi/WebPlotDigitizer. Then, using these size structures, we used the fully size-dependent transmission model to estimate the probability of successful infection by a schistosome given that schistosome makes irreversible contact with a snail in the population for each time point. We took this approach due to the lack of data on waterbody size in this dataset, and therefore we were unable to estimate snail infection prevalence since snail density was unknown. However, the availability of snail abundance and body size data allowed us to take the point of view of a parasite encountering the waterbody and confronted with different snail size structures at different points in time. Specifically, to estimate probability of a parasite succeeding in infecting a snail, for each size class (1 mm increment) of snail we estimated the probability that the schistosome miracidium would contact and subsequently infect a snail in that single size class based on the relative abundance of snails in that size class and the size-dependent exposure and susceptibility traits of all snails in the population at that time. We then summed infection probabilities across all snail size classes for each time point to get an overall probability of infection (Eq. 12):

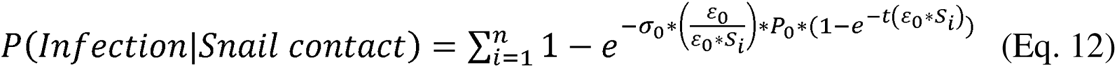

**Figure 1.**
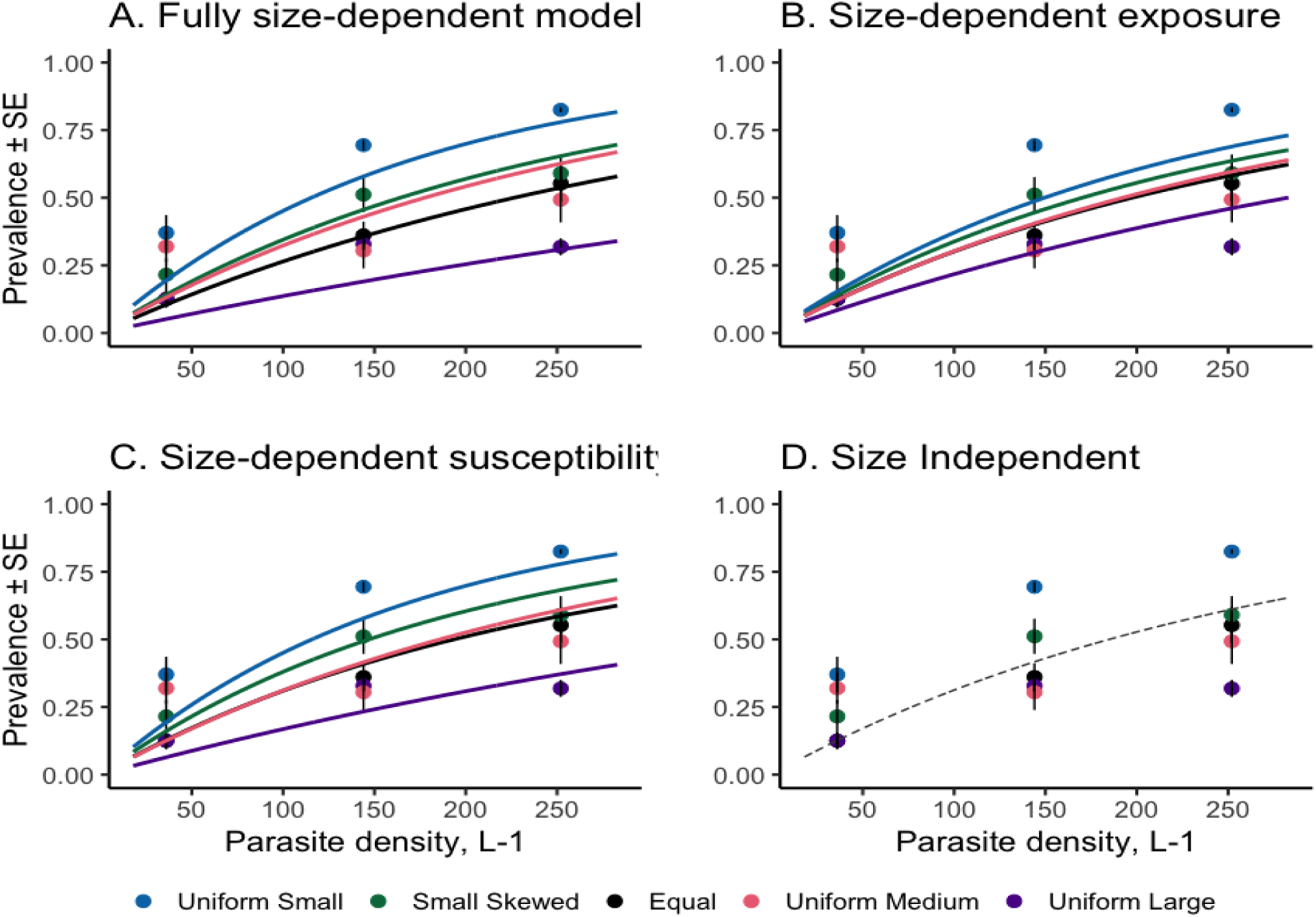
Schistosome prevalence in snail populations with differing size structures. Points represent mean ± SE prevalence in each population structure and lines represent best model fits. Each panel depicts one of the candidate transmission models; experimental data is identical between panels. In our transmission trials, uniform small populations had a significantly greater prevalence than the uniform large uniform medium and equal size structures, and prevalence in the uniform large size structure was significantly less than the small skewed and the uniform small. A fully size-dependent transmission model (**A**) in which exposure and susceptibility vary with body size outperformed all other candidate models. Allowing only exposure to vary with body size (**B**) tended to underestimate prevalence in populations with a greater proportion of small individuals, while a model in which only susceptibility varies with body size (**C**) performed better than exposure-dependence, it did especially poorly at predicting prevalence of uniformly large populations. A null model with no size-dependence (**D**) is unable to make different predictions in differently size-structured populations, and thus has a very poor fit to the observed data.

We estimated 95% confidence intervals for this quantity by drawing estimated transmission parameters from the joint 95% confidence interval set inferred from the experimental results (Bolker 2008). We contrasted these estimates to the 95% confidence interval for infection success given exposure for the size-independent (null) model, which makes a constant prediction of parasite success, independent of size structure.

## III. Results

### Experimental Transmission Experiments

Population prevalence for snails varied by size structure and parasite density (Fig. 1). The uniform small size structure had the greatest prevalence at each parasite density (36 parasites: 0.37 ± 0.026 [mean ± SE]; 144 parasites: 0.69 ± 0.078; 256 parasites: 0.83 ± 0.034) and this difference was significantly greater than the uniform large (z ratio = -5.425, p<.0001), uniform medium (z ratio = -3.868, p=0.0010) and equal size structures (z ratio = -4.203, p= 0.0003). The uniform large size structure had the smallest prevalence of the tanks at the lowest and highest parasite densities (36 parasites: 0.13 ± 0.011; 256 parasites: 0.32 ± 0.13), and prevalence for the uniform large size structure was significantly less than the small skewed (z ratio = 3.480, p= 0.0046) and the uniform small (z ratio = -5.425, p <.0001).

Within size classes, the average prevalence for small snails varied by population size structure and parasite density (Fig 2). Regardless of population size structure, large snails were less likely to be infected than small snails (z ratio = -3.67, p=0.0022). Small snails in the uniform small size structure had the highest prevalence across parasite densities (Fig 2.A), and were significantly more likely to be infected than small snails in the equal size structure treatment (z ratio = -5.102, p<.0001). Small snails in the small skewed size structure were also significantly more likely to be infected than small snails in the equal size structure (z ratio = -3.552, p = 0.0011). Medium and large snails did not show significant differences in infection between size structures (Fig 2.B,C).

**Figure 2.**
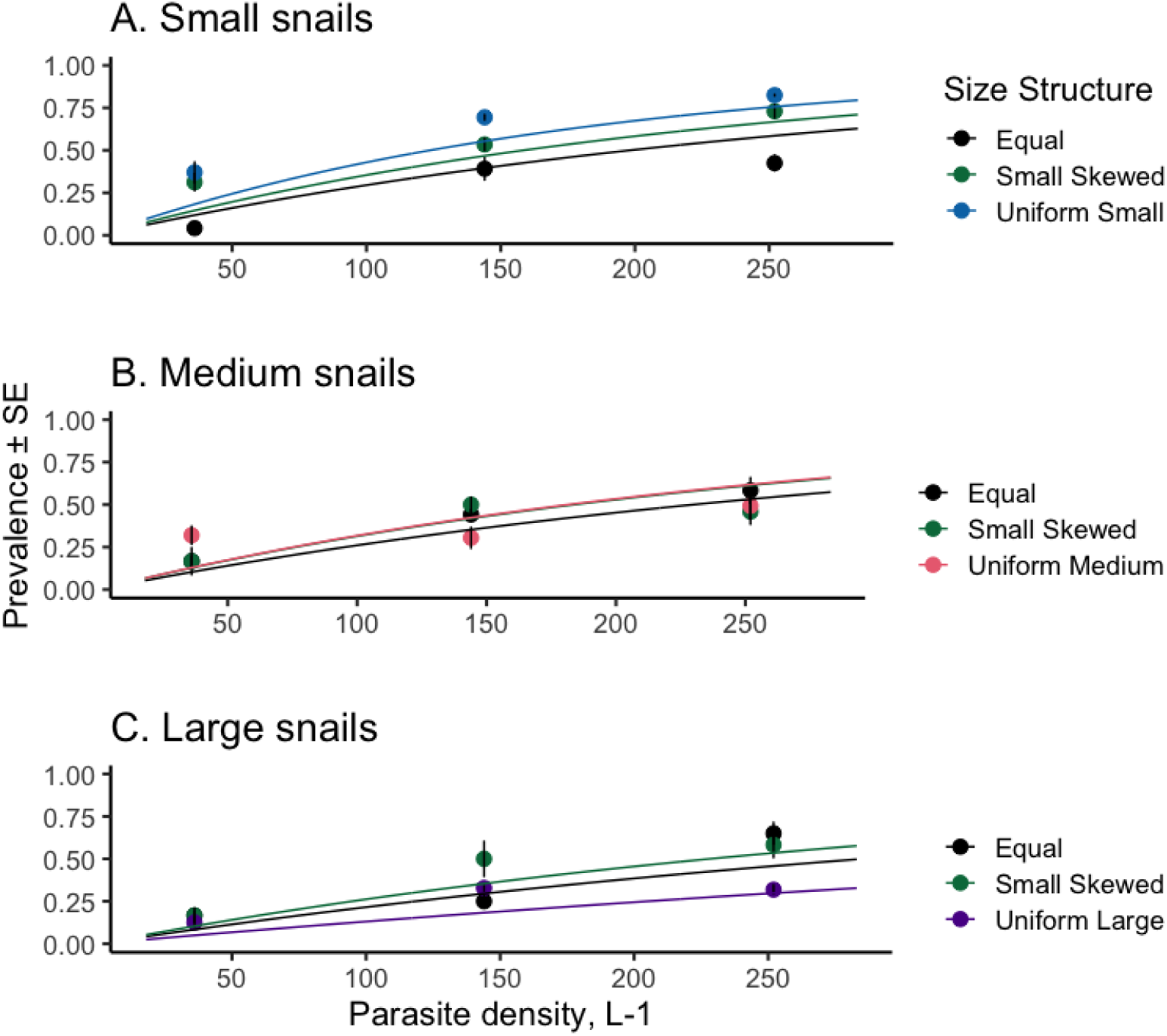
Schistosome prevalence within size classes in snail populations with differing size structures. Points represent average prevalence in each population structure, n=4 replicates per treatment; lines represent fits from the winning candidate model with full size dependence. A) Small snails B) Medium snails C) Large snails. Size structures with a greater proportion of large conspecifics resulted in lower infection prevalence in the small snails within those populations. Medium and large snails did not show significant differences in infection between size structures.

### Transmission model competition

The size-independent null model performed poorly (Table 1). It either over or underestimated transmission depending on the population size-structure (Fig 1D). Exposure-only size dependence and susceptibility-only size dependence performed better (Table 1), but still tended to underestimate schistosome prevalence especially at lower parasite densities (Fig 1B, 1C). A fully size-dependent model performed far better than other candidate models (Table 1), despite similar issues in underestimation at the lowest parasite density. An exponential relationship between body size and exposure and susceptibility outperformed a linear relationship (Appendix S1:Figure S1, Appendix S1:Table S1), therefore we focus in the main text on comparing size-based models with exponential function(s) to the null model. Maximum likelihood estimates for size-dependent exposure and susceptibility from the fully size-dependent model allow for predictions of exposure(ε), susceptibility(σ), and transmission (β = ε x σ) based on body size (Fig 3). Experimental data on size-based susceptibility correspond well to these predictions (Fig 3B) and the transmission parameter is more strongly influenced by susceptibility, resulting in a prediction of overall lower transmission with increasing body size (Fig 3C).

**Figure 3.**
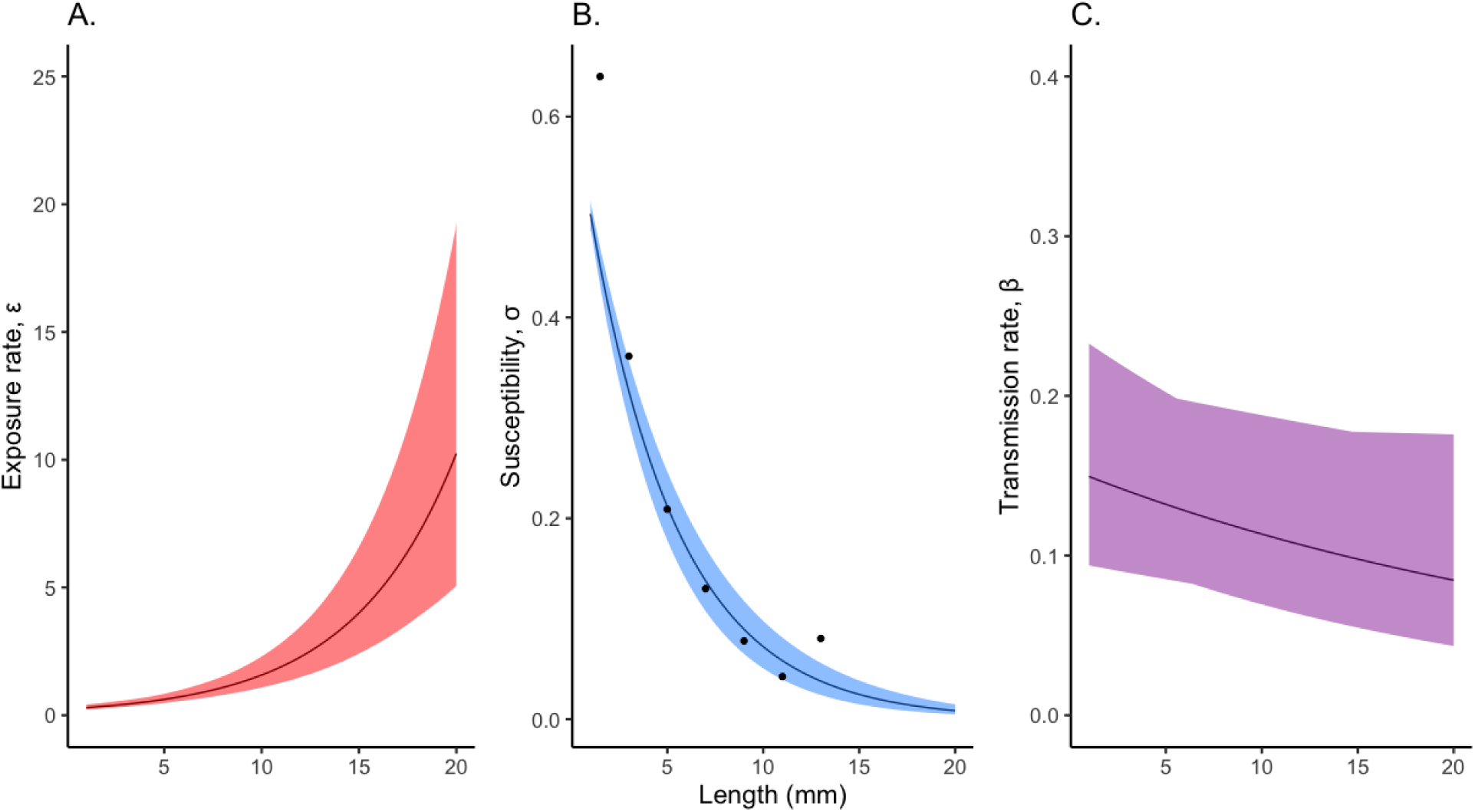
Estimates for exposure, ε(L), (A), susceptibility,σ(L), (B), and overall transmission rate, β(L) = ε(L)*σ(L) (C) as functions of snail length. Lines represent the maximum likelihood of estimated parameters across body size of snails. Shading represents the minimum and maximum from the 95% confidence interval of parameters estimated. Points in (B) are from experimentally observed schistosome susceptibility across snail sizes from individual exposures, while the other two parameters are inferred from the full dataset.

**Figure 4.**
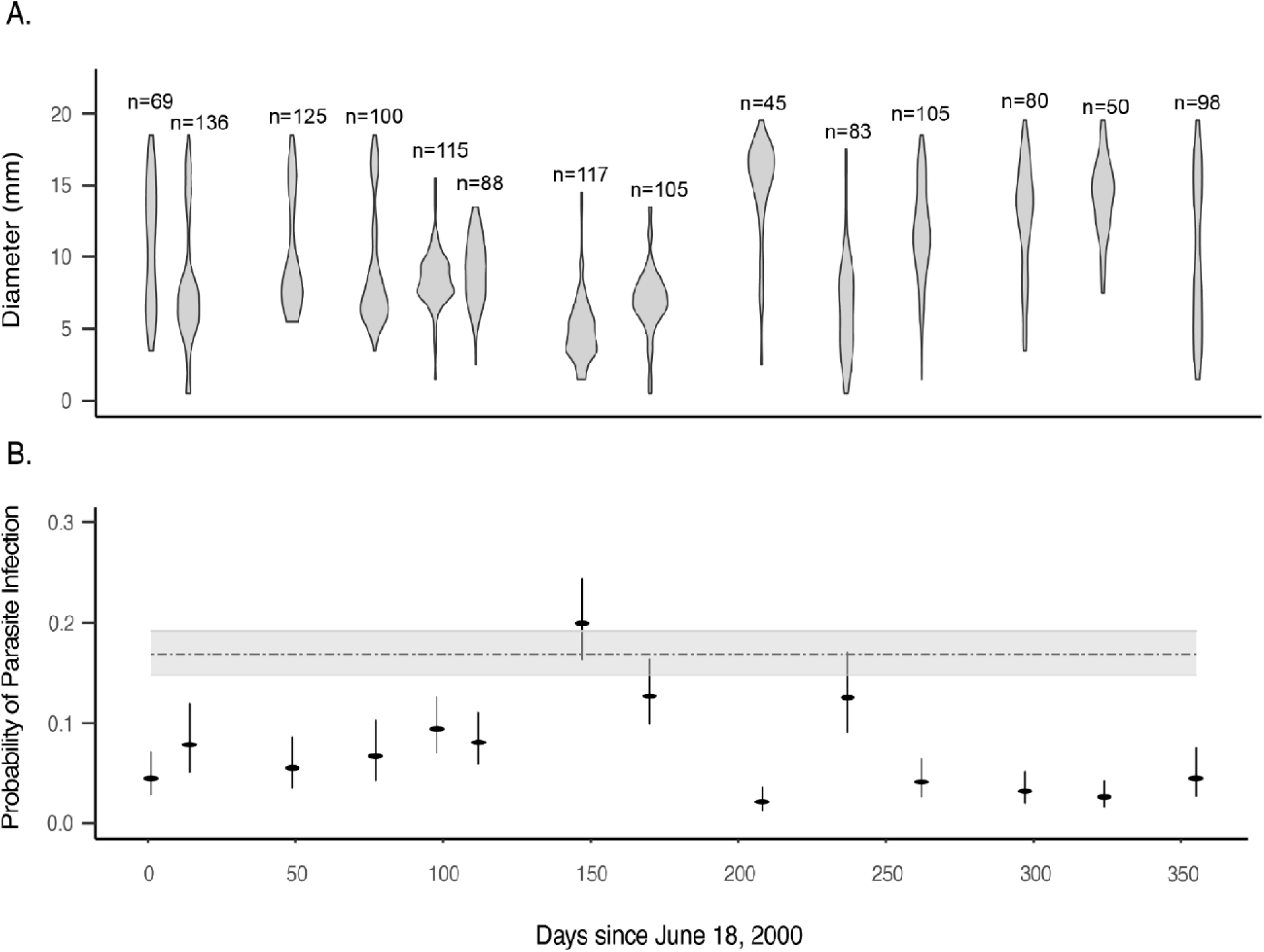
Predicted infection success of a single schistosome parasite given exposure of a snail in a natural population with varying body size structure over time. A) Distribution of body sizes from a population of *Biomphalaria tenagophilia* in del Plata Basin, Argentina from 2000-2001(Rumi et al. 2009). Number of snails sampled at each timepoint represented as n=. B) Points are the mean ± 95% confidence interval of the predicted infection success of a single schistosome introduced into this snail population from simulated transmission trials using the fully size-dependent model. Grey line is the predicted successful infection from a transmission model that ignores size-based exposure and susceptibility. Substantial changes in snail population size structure cause month-to-month estimates of schistosome success to diverge substantially from predictions that ignore size dependence. In particular, the size-null prediction tends to overestimate schistosome success except at the times in which the population is heavily dominated by small snails.

**Table 1.**
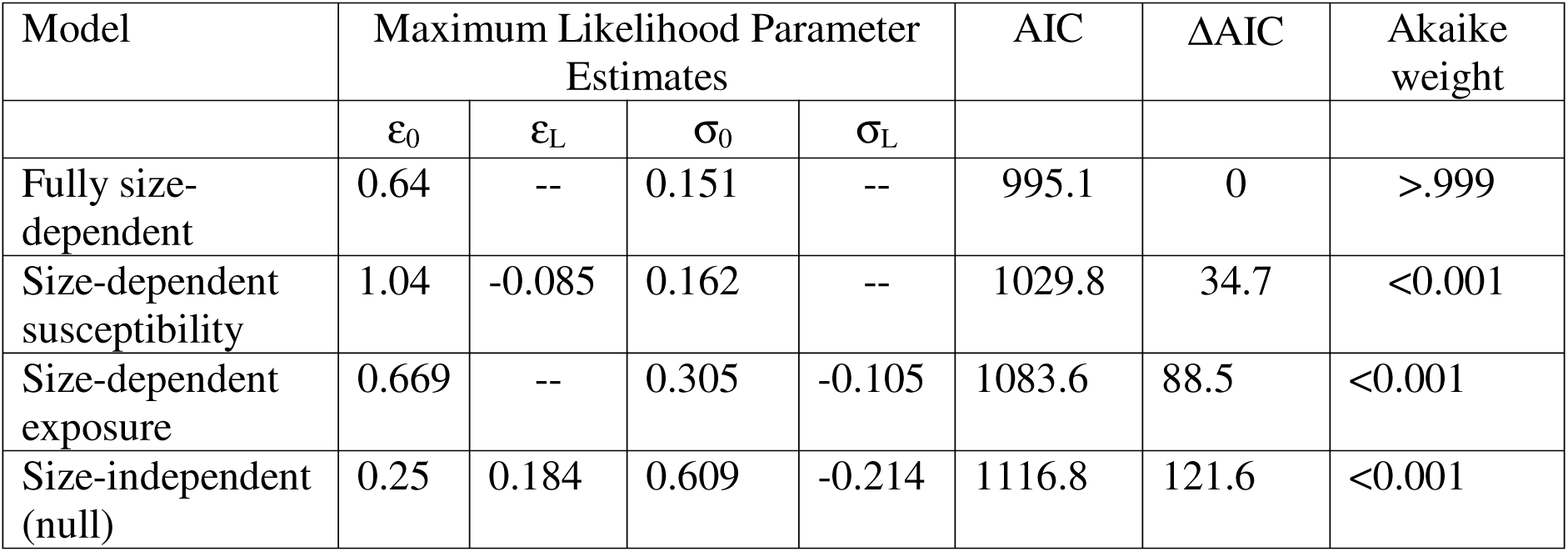
Parameter estimates and model selection statistics.

| Model | Maximum Likelihood Parameter Estimates | | | | AIC | ΔAIC | Akaike weight |
| --- | --- | --- | --- | --- | --- | --- | --- |
| | $\epsilon_0$ | $\epsilon_L$ | $\sigma_0$ | $\sigma_L$ | | | |
| Fully size-dependent | 0.64 | -- | 0.151 | -- | 995.1 | 0 | >.999 |
| Size-dependent susceptibility | 1.04 | -0.085 | 0.162 | -- | 1029.8 | 34.7 | <0.001 |
| Size-dependent exposure | 0.669 | -- | 0.305 | -0.105 | 1083.6 | 88.5 | <0.001 |
| Size-independent (null) | 0.25 | 0.184 | 0.609 | -0.214 | 1116.8 | 121.6 | <0.001 |

### Implications for *S. mansoni* dynamics

The size structure of a natural *Biomphalaria* population varied substantially during the year (Fig 4A). Using these size structure data and our winning fully size-dependent model, we estimated that the probability of establishing a successful infection for a single schistosome miracidium encountering a snail in this population would also vary over the course of the year due to the influence of snail body size on transmission. We took this approach of generating predictions from our model due to the lack of available data from field settings that includes both snail body size and infection prevalence. When small snails are overrepresented in the population, e.g. day 150, the predicted success of a schistosome is much higher than when the population is skewed towards larger individuals, e.g. day 208 (Fig 4B). Importantly, the null and size-dependent models yielded similar estimates in only 3 of 14 months. Overall, the size-dependent model predicted lower parasite success than the null model, suggesting that ignoring size structure could lead to overestimation of transmission potential.

## Discussion

Natural populations are heterogenous, and in order to create ecological theory that can reflect observed dynamics or predict future outcomes we must find tenable ways to address this heterogeneity (Bolnick et al. 2011). It is particularly important to include relevant heterogeneity in transmission, because this process shapes the ecological and evolutionary trajectory of disease dynamics (McCallum et al. 2001). Transmission models that do not incorporate heterogenous host traits often fit data poorly (Dwyer et al. 2010), compromising their applicability to natural populations. Therefore, identifying the traits that drive transmission and the directionality of their effects can improve theoretical predictions for natural dynamics and better assess interventions.

However, we do not suggest that disease ecologists embark on an endless hunt to characterize all heterogenous host traits. Instead, focusing on traits that either integrate numerous aspects of a host’s ecology, have the potential to regulate multiple downstream processes impacting transmission, or both, can greatly reduce unexplained variability while still being tenable to include mathematically and empirically. Body size is trait that satisfies both of these criteria for many disease systems, as it varies amongst individuals due to many processes such as age and nutrition (Kooijman, S.A.L.M. 2010), and it profoundly impacts many ecological processes such as predation, resource competition, and reproductive success(Wellborn 1994, Hopcraft et al.

2010). This leads to its utility as a tractable trait to include in transmission models because often an individual’s history resulting in their current body size is usually impossible to reconstruct for wild organisms. Further exploring the role of body size in other host-parasite systems may reveal correlations and trade-offs between host physiology, behavior, and transmission.

For decades, it has been known that larger *Biomphalaria* snails experience a higher exposure to schistosomes but a lower susceptibility to infection (Niemann and Lewis 1990). However, the relevance of this phenomenon to epidemiological dynamics has not been determined. Here we demonstrate the epidemiological relevance of parasite transmission in size-structured populations. We showed that this individual-level phenomenon can have population-level consequences for schistosome prevalence: differently size-structured populations experienced very different schistosome transmission dynamics in our experimental trials. Populations with a greater proportion of large sized snails had lower schistosome prevalences, while populations of uniformly small snails experienced the greatest prevalence. From the perspective of an individual snail, the body size of immediate neighbors is also important for transmission: small snails had lower prevalence of schistosomes when in populations with a greater proportion of large conspecifics. This observation supports the hypothesis that individuals with relatively high exposure rates to parasites can shield their conspecifics from infection, a form of transmission interference akin to the much-discussed dilution effect that is typically considered in multi-species host communities (Keesing et al. 2006, Shaw and Civitello 2021). While our experimental design only captured short-term transmission dynamics, explicitly incorporating host body size into schistosome transmission models will allow for future extensions of this work to explore longer term outcomes and include ecological interactions that can impact body size such as resource competition (Malishev and Civitello 2020). Furthermore, this size-explicit transmission model can now be incorporated into models that evaluate the entire schistosome lifecycle, and better predict transmission risk to humans based both on the impact of snail size at the time of transmission to the snail and also snail production of cercariae, the life stage which is infectious to people, because cercarial production can be size and resource dependent (Civitello et al. 2022).

Incorporating size structure into schistosome transmission models could also improve our understanding of seasonal variation in schistosome transmission to intermediate snail hosts. We found that for a well-characterized natural *B. glabrata* population, temporal variation in size structure was large enough to cause substantial variation in predicted transmission success in snails for schistosomes. It’s important to note that this is not an empirical result: it is using our mechanistic model to generate a hypothesis on the impact of size-structure on schistosome dynamics in endemic settings. To our knowledge, no available public datasets contain the necessary data on snail density, body size, and infection prevalence to directly compare to our model predictions. Our hope is that the stark predicted difference in parasite infection success between the null and the fully size-dependent model will motivate future work to collect the necessary data to directly test this hypothesis. Numerous biotic and abiotic factors, such as predation, resource competition, temperature, and nutrient availability, can impact host snail growth and therefore population size structure across space and time. Furthermore, schistosome intervention efforts can be costly and labor-intensive, but characterizing the seasonal patterns in size structure may allow for more targeted approaches at times when the chance of transmission is highest, such as following a large reproductive burst when the population has a large proportion of neonates.

Hosts are not identical, despite the plethora of transmission models that treat them as such. Scaling up the effects of heterogenous host traits on transmission to investigate population-level effects can allow for better inference and prediction, especially when using traits such as body size that are measurable and can reflect correlations between numerous ecological processes. Our work shows the importance of this approach in the snail-schistosome system. Tackling the influence of host heterogeneity in transmission may be key for resolving many unanswered questions in disease ecology.

## Acknowledgements

We thank members of the Civitello Lab for their comments on this manuscript. KS was supported by the National Institute of Allergy and Infectious Diseases of the National Institutes of Health under Award Number T32AI138952 and Award Number F31AI147611. DJC was supported by NIH 1R01 AI150774-01. The content is solely the responsibility of the authors and does not necessarily represent the official views of the National Institutes of Health.

## Author Contributions

KES and DJC conceived and designed the study. KES, REC, RS performed experiments. KES, REC, DJC analyzed data. KES and REC wrote the manuscript with input from DJC.

#### Box 1.

**Model construction**

We constructed deterministic transmission models that tracked changes in the densities of susceptible hosts, S_i_ and infected hosts, I_i_, in an arbitrary number of host size classes, *i*, as well as free-living parasites, P, with differential equations:

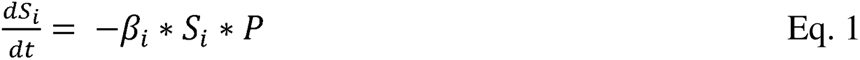

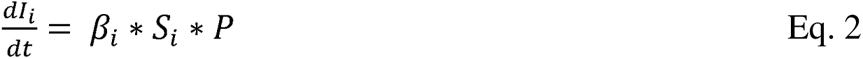

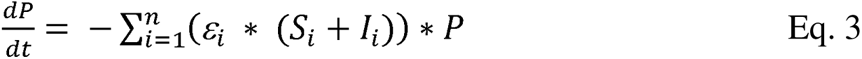

We parameterized these models based on a short-term infection experiment, therefore we ignored processes that occur at longer time scales such as birth and deaths of hosts and parasites (Civitello and Rohr 2014). For each model we broke down the transmission rate (β*_i_*) into two main components, exposure (ε*_i_*) and susceptibility (σ*_i_*), that we allowed to vary with size (L *_i_*):

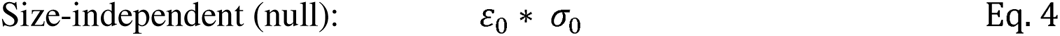

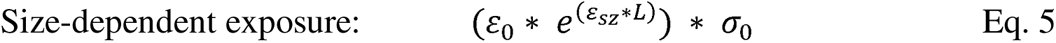

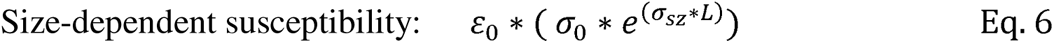

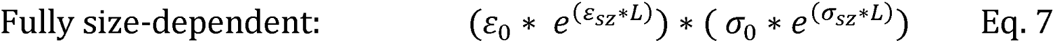

Prevalence can be directly calculated within each size class (i), based on the initial density of susceptible individuals in each size class, S(0)_i_, :

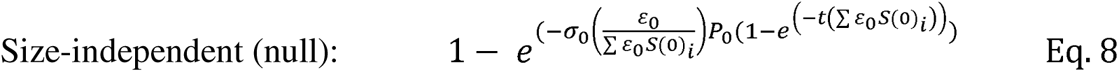

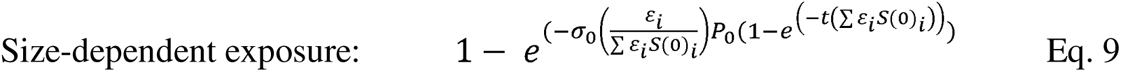

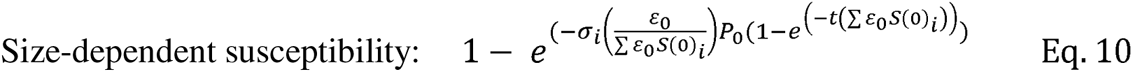

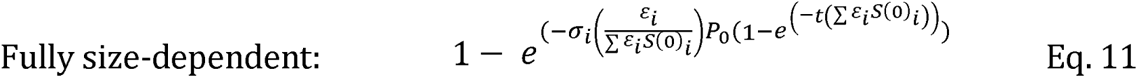

