## Supplementary Material for "Parasite transmission in size-structured populations"

**Table of contents:**

- Table S1
- Figure S1

**Table S1**. Model competition of linear and exponential size-based parameters

| Model | Transmission rate | AIC | ΔAIC | Akaike weight |
| --- | --- | --- | --- | --- |
| Exponential size-based relationship | ${(\varepsilon}_{0}* e^{\left( \varepsilon_{sz}*L \right)})*( \sigma_{0}*e^{\left( \sigma_{sz}*L \right)})$ | 995.1 | 0 | 1 |
| Linear size-based relationship | ${(\varepsilon}_{0}+ \varepsilon_{sz}*L)* {(\sigma}_{0}+ \sigma_{sz}*L)$ | 1078.2 | 83 | <0.001 |

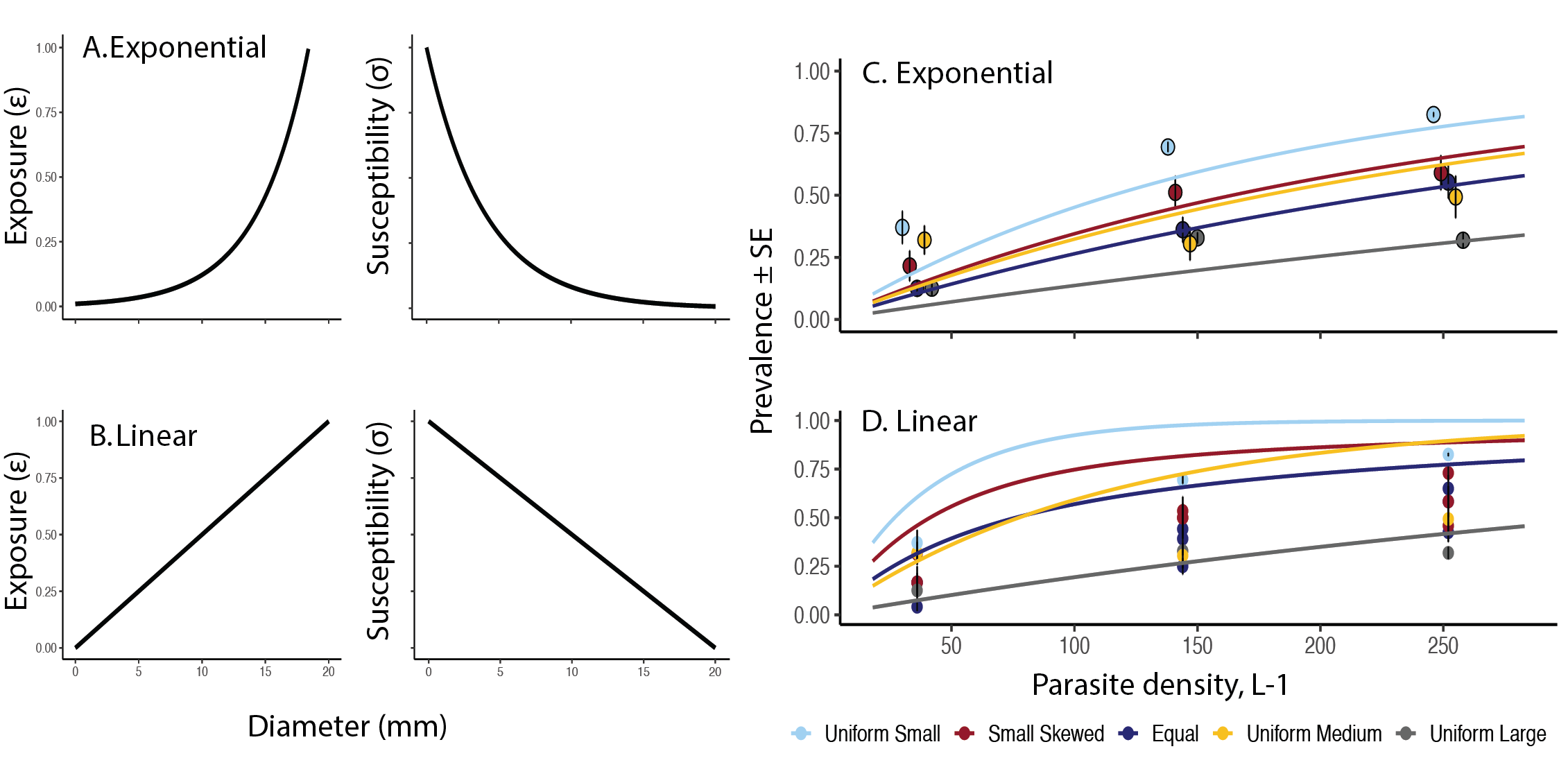

**Figure S1.** A) An illustration of an exponential relationship between body size and exposure and body size and susceptibility. B) An illustration of a linear relationship between body size and exposure and body size and susceptibility. C) Fitting a fully size-dependent model to data based on an exponential relationship between body size, exposure, and susceptibility. Identical to main text Figure 1A. D) Fitting a fully size-dependent model to data based on a linear relationship between body size, exposure, and susceptibility. A linear relationship for these parameters has a very poor fit and was not considered in further analyses.
